# Expression of sex steroid receptors and sex differences of Otp glutamatergic neurons of the medial extended amygdala

**DOI:** 10.1101/2024.10.23.619216

**Authors:** Alba González-Alonso, Lorena Morales, Elisenda Sanz, Loreta Medina, Ester Desfilis

## Abstract

The medial extended amygdala (EAme) is part of the social behavior network and its subdivisions show expression of sex steroid receptors, which participate in the regulation of sexually dimorphic behaviors. However, EAme subdivisions are highly heterogeneous in terms of neuron subtypes, with different subpopulations being involved in regulation of different aspects of social and non-social behaviors. To further understand the role of the different EAme neurons and their contribution to sexual differences, here we studied one of its major subtypes of glutamatergic neurons, those derived from the telencephalon-opto-hypothalamic domain that coexpress *Otp* and *Foxg1* genes during development. Our results showed that the vast majority of the Otp glutamatergic neurons of the medial amygdala and BSTM in both sexes express *Ar*, *Esr1 (ERα)* and *Esr2 (ERβ*) mRNA. Moreover, the high percentage of receptors expression in the Otp neurons (between 93 and 100%) indicates that probably the majority of the Otp neurons of EAme are coexpressing the three receptors. In addition, Otp neurons of the posterodorsal medial amygdala have a larger soma and occupy more space in males than in females. These and other features of the Otp neurons regarding their expression of sex steroid receptors likely contribute to some of the sexually-dimorphic behaviors regulated by EAme.

## Introduction

The medial extended amygdala (EAme) is an important associative nuclear complex that is part of the social behavior network, a brain functional system composed by areas and nuclei that are essential for social cognition (social recognition, discrimination and affiliation) and are involved in a variety of social behaviors, like parental, sexual or agonistic (Newman, 1999; Rasia-Filho et al., 2012; Medina et al., 2019; Raam and Hong, 2021; Lischinsky et al., 2023; McCarthy, 2023). As described by Newman in 1999, this network is composed of the EAme (including the medial amygdala and the medial bed nucleus of the stria terminalis [BSTM]), the lateral septum (LS), the medial part of the preoptic area (POA), the anterior hypothalamus (AH), the ventromedial hypothalamus (VMH), the periaqueductal gray (PAG) and the adjacent tegmentum. Additional centers, like the paraventricular hypothalamic nucleus (PVN) and the dorsomedial hypothalamus (DMH), could also be considered part of the network because of their relevance in regulating neuroendocrine and/or other aspects of social behavior (Goodson and Kingsbury, 2013; Chen and Hong, 2018; DiMicco at al., 2002; Kataoka et al, 2020). The centers of this system are connected with each other and have a strong expression of sex hormone receptors, which regulate the activity of this network (Simerly et al., 1990; Shughrue et al., 1998; Newman, 1999; Goodson and Kingsbury, 2013; Chen and Hong, 2018).

Many aspects of social behavior show strong sexual differences (Yang and Shah, 2014; Chen and Hong, 2018; Fang et al., 2024), which start to develop by the differences in availability of sex steroids during prenatal and perinatal periods, and the interactions of the circulating sex hormones with different neuron subpopulations of the brain social behavior network that express specific receptors (Han and De Vries, 2003; Arnold, 2009; McCarthy and Arnold, 2011; McCarthy, 2023). In males, testosterone produced during the prenatal and perinatal periods and transformed to estradiol by aromatase masculinizes the brain, organizing the differentiation of neural circuitries involved in sex specific social behaviors (De Vries and Miller, 1998; Han and De Vries, 2003; De Vries and Panzica, 2006). This could underlie differences between sexes in the prevalence of some neurodevelopmental disorders with social behavior components, such as autism spectrum disorder, with higher prevalence in males (Newman, 1999; Cooke and Woolley, 2005; Morris et al., 2008; Rasia-Filho et al., 2012; Yang and Shah, 2016). The differences become more pronounced from puberty, with the sexually dimorphic production of androgens, estrogens, and progesterone (Arnold, 2009).

Regarding the EAme, studies in different species including rodents have shown that this nuclear complex shows anatomical differences, with differences in volume, neuron number, size and complexity (Cooke and Woolley, 2005; Morris et al., 2008; Rasia-Filho et al., 2012). These differences are observed specially in the posterodorsal subnucleus of the medial amygdala (MePD) and in the BSTM, which are larger in males than in females both in rats and mice, and have bigger cells (Segovia and Guillamón, 1993; Cooke and Woolley, 2005; Cooke et al., 2007; Morris et al., 2005, 2008; Rasia-Filho et al., 2012). Unlike the rat, there are no differences between sexes in the total number of neurons in the mouse MePD (Cooke and Woolley, 2005; Morris et al., 2005, 2008; Rasia-Filho et al., 2012). In addition, differences in the expression of different neuropeptides have also been observed; for example, in the EAme (MePD and/or intraamygdaloid BST, plus BSTM) of rats, mice and other rodents, the expression of vasopressin, substance P or cholecystokinin is more intense in males than in females (De Vries and Miller, 1998; Cooke and Woolley, 2005; Otero-García et al., 2014). In contrast, oxytocin cells are more abundant in PVN of females compared to males (Xu et al., 2010), and the expression level of oxytocin in the female hypothalamus is also higher (Häussler et al., 1990). Vasopressinergic and oxytocinergic subsystems play key roles in the expression of sexually dimorphic behaviors and show variations between species at the level of peptide and/or peptide receptor expressions that correlate with sociality features, such as social affiliation and pair bonding (Insel and Shapiro, 1992; Insel et al., 1994; Donaldson and Young, 2008; Xu et al., 2010). Notably, the brain expression differences and functions of these subsystems are influenced by the levels of gonadal hormones and their receptors (De Vries et al., 1983; De Vries and Miller, 1998; Caldwell and Moe, 1999; De Vries and Panzica, 2006).

Sex differences in the brain expression of sex steroid receptors are variable depending on the species, the developmental stage of the animals and if the molecule analyzed is the mRNA or the protein (Lu et al., 1998; Równiak, 2017; Cooper et al., 2021; Cara et al., 2021). Recent research has shown that in mice such differences of expression are subtle and easier to detect when studying distinct molecularly defined neuron subpopulations (Shah et al., 2004; Pfau et al., 2023). This is particularly important in the EAme, as this center is highly heterogenous, with a wide variety of neuronal subtypes that originate in distinct embryonic domains, have different gene expression signatures, and are involved in different functional pathways (García-López et al., 2008; Hirata et al., 2009; Carney et al., 2010; García-Moreno et al., 2010; Bupesh et al., 2011; Lischinsky et al., 2017, 2023; Ruiz-Reig et al., 2017; Morales et al., 2021; reviewed by Medina et al., 2011, 2017, 2023; Abellán et al., 2013). Some of these neuron subtypes originate in the subpallium and are GABAergic (García-López et al., 2008; Hirata et al., 2009; Carney et al., 2010; Sokolowski and Corbin, 2012; Lischinsky et al., 2017; Medina et al., 2023), but there is also an important number of glutamatergic neurons with different embryonic origins from telencephalic and extratelencephalic domains (reviewed by Abellán et al., 2013; Medina et al., 2023), and recent studies have shown that, in some subdomains of EAme nuclei, they are as numerous as the GABAergic ones (García-Moreno et al. 2010; Morales et al., 2021). While the activation of different subsets of GABAergic neurons promotes different types of social interactions (Sokolowski and Corbin, 2012; Lischinsky et al., 2017, 2023), the role of the glutamatergic neurons is less clear and may differ depending on the embryonic origin (Abellán et al., 2013) and/or EAme subdivision (Choi et al., 2005). Regarding glutamatergic neurons of MePD, their activation appears to inhibit social interactions and promote self-grooming, which resembles autistic-like behaviors (Hong et al., 2014; Yang and Shah, 2016; Johnson et al., 2021; Kwon et al., 2021). This shows that the glutamatergic neurons also seem to play an important role in the regulation of behavior, although the mechanisms for this regulation remain unknown. One of the most important and numerous glutamatergic neurons of MePD, as well as many of those in other EAme subdivisions, are the neurons that during development express the homeobox protein orthopedia (OTP). These were thought to originate in the supraopto-paraventricular embryonic domain (SPV) of the alar hypothalamus (Bardet et al., 2008; García-Moreno et al., 2010). However, recently, our group found that in mouse the majority of these neurons coexpresses the transcription factors FoxG1 (typical of telencephalon) and OTP, and originate in a new radial embryonic domain near the frontier between subpallium and hypothalamus, called the telencephalon-opto-hypothalamic domain or TOH (Morales et al., 2021). The SPV core also produces Otp neurons that can be distinguished from the TOH neurons because they do not express Foxg1. In mouse, OTP is critical for the differentiation of forebrain vasopressinergic and oxytocinergic neurons (Wang and Lufkin, 2000), but it is likely that these cells are produced in SPV core, and not the TOH (Morales et al., 2021). The majority of SPV core-derived neurons are found in the PVN, specially its central and ventral parts, and in the adjacent hypothalamus, but only a few of them migrate to the EAme (Morales et al., 2021; Medina et al., 2023). Although MePD and BSTM are sexually dimorphic, it is unclear if their glutamatergic cells contribute to or participate in such differences. Our aim was to investigate if the Otp glutamatergic neurons of the EAme present sexual differences in the volume they occupy, their cell number and their expression of gonadal hormones’ receptors. To solve the problem of the downregulation of *Otp* after development, that may hinder our ability to identify all the Otp neurons in adult animals, we used the *Otp-eGFP* transgenic mouse line. This line has already been characterized and found appropriate as GFP recapitulates *Otp* expression during embryonic development in the amygdala and hypothalamus (Morales et al., 2021, 2022). Brains of *Otp-eGFP* mice of both sexes were employed to study morphological details of the Otp neurons and their coexpression of sex-steroid receptors by combining immunofluorescence with fluorescent *in situ* hybridization. The areas studied were the different subdivisions of the EAme (subdivisions of the medial amygdala and BSTM), all of them with a high density of Otp neurons, the majority of which derive from the TOH. We also analyzed other centers of the social behavior network, which contain Otp cells of different embryonic origins (such as PVN) and/or display abundant Otp terminals of intrinsic and/or extrinsic origins, such as LS, DMH, VMH, and PAG (Morales et al., 2021).

## Material and Methods

### Animals

In this study we used *Otp-eGFP* transgenic mice (*Mus musculus*, Tg (Otp-EGFP) OI121Gsat/Mmucd; Mutant Mouse Resource & Research Centers, MMRRC supported by NIH, University of California at Davis, USA) with permanent labeling of Otp neurons and their projections (Morales et al., 2021, 2022). The *Otp-eGFP* transgenic mice was validated previously for studies of Otp neurons of EAme and hypothalamus (Morales et al., 2021, 2022).

Progenitors and weaned-off postnatal animals were housed in groups of three to five at 22 ± 2°C on a 12-hour light/dark cycle, with food and water ad libitum, in the rodent animal facility of the University of Lleida (REGA license no. ES251200037660). All the animals were treated according to the regulations and laws of the European Union (Directive 2010/63/EU) and the Spanish Government (Royal Decrees 53/2013 and 118/2021) for the care and handling of animals in research. All the protocols used were approved by the Committees of Ethics for Animal Experimentation and Biosecurity of the University of Lleida.

### Expression of sex steroid receptors mRNA in Otp neurons

For expression of sex steroid receptors, we used 16 Otp-eGFP animals of both sexes, aged as follows: 7 pre-weaned (P19), 4 mature adults (P60), and 5 middle-aged animals (P174-P271). For conventional *in situ* hybridization, we used 10 animals, 5 males and 5 females. For fluorescent *in situ* hybridization combined with immunofluorescence, we used 7 animals, 3 males and 4 females (one of the animals was used for both procedures).

### Tissue collection and fixation

The animals were deeply anesthetized with sodium pentobarbital (0.1 mg/g; i.p.) and then transcardially perfused with 0.9% saline solution (0.9% NaCl), followed by phosphate-buffered (PB; 0.1M, pH 7.4) 4% paraformaldehyde (PFA). After dissection, the brains were postfixed by immersion in 4% PFA overnight at 4°C.

### Sample preparation and sectioning

The brains were embedded in 4% low-melt agarose (LOW EEO, Laboratorios Conda S.A., Spain) dissolved in 0.1M phosphate-buffered saline (PBS) and sectioned in 80μm thick frontal sections using a vibratome (Leica VT 1000S; Leica Microsystems GmbH, Germany). The sections were collected in 4°C PBS and then processed for conventional *in situ* hybridization or fluorescent *in situ* hybridization combined with single immunofluorescence.

### Conventional *in situ* hybridization

Brain sections were processed for *in situ* hybridization using digoxigenin-labelled riboprobes produced following the procedure previously described by Abellán et al., 2014. The antisense riboprobes labelled with digoxigenin were synthesized using Roche Diagnostics (Germany) protocols from cDNAs of the genes of interest: *Ar*, *Esr1* (ERα), and *Esr2* (ERβ) (Table 1).

**Table 1.** Mouse genes for riboprobes.

| Mouse genes for riboprobes |  |  |  |
| --- | --- | --- | --- |
| Gene | Insert size | Purchased to | GenBank accession no. |
| *Ar* | 435 pb | Source Bioscience (EST Clone: IMAGp998F183G15Q) | AI182177 |
| *Esr1* (alpha) | 593 pb | Source Bioscience (EST Clone: IMAGp998A121081Q) | AA023625 |
| *Esr2* (beta) | 463 pb | Source Bioscience (EST Clone: IMGSp981B08229D) | BX630139 |

Before the hybridization, the tissue was permeabilized using 0.1% Tween-20 in PBS (PBT; 0.1 M) and washed three times for 10 minutes each. After this, the sections were prehybridized in hybridization buffer (HB) containing 50% deionized formamide (Sigma-Aldrich), 6.5% standard saline citrate (pH 5; 2M), 1% ethylenediaminetetraacetic acid (EDTA; pH 8; 5M; Sigma-Aldrich), 0.2% Tween-20, 1 mg/ml of yeast tRNA (Sigma-Aldrich), 100 μg/ml of heparin (Sigma-Aldrich), completed with RNase and DNase free water (Sigma-Aldrich). The prehybridization lasted at least 2-4 hours at 58°C. Then the sections were hybridized overnight at 58°C in HB containing 0.5-1 μg/ml of the corresponding riboprobe. The next day the hybridized sections were washed first in HB at 58°C, then in a 1:1 mix of HB and MABT (maleic acid buffer containing Tween 20; 1.2% maleic acid, 0.8% NaOH, 0.84% NaCl, and 0.1% Tween-20) at 58°C and, after this, in MABT at room temperature with gentle stirring. Then, to avoid unspecific binding, the sections were blocked using a solution containing 20% blocking reagent BBR and 20% sheep serum in MABT for 2 hours at room temperature. Following this, the sections were incubated overnight at 4°C with a sheep anti-digoxigenin antibody conjugated with alkaline phosphatase (Roche Diagnostics) diluted 1:3500 in MABT. After washing in the same buffer and in B2 buffer (100 mM Tris–HCl, 100 mM NaCl, 50 mM MgCl2, pH 7.5), the sections were finally revealed with NBT (nitroblue tetrazolium) and BCIP (5-bromo-4-chloro-3-indolyl phosphate) (both from Roche Diagnostics) mixed in B2, and then rinsed, mounted and coverslipped with Permount.

### Fluorescent *in situ* hybridization combined with immunofluorescence

Some series of brain sections of *Otp-eGFP* mice were processed for fluorescent *in situ* hybridization (FISH) using an indirect method. For this procedure, we used the same antisense riboprobes mentioned above to detect the mRNA of sex steroid receptors (Table 1). The FISH was followed by immunofluorescence for GFP to see if the Otp neurons of selected brain regions express the receptors.

Before starting, the tissue was permeabilized using 0.1% Tween-20 in PBS (PBT; 0.1 M) and washed three times for 10 minutes each. After this, the sections were prehybridized in hybridization buffer (HB) (prepared as explained above). The prehybridization lasted 2-4 hours at 58°C. Then the sections were hybridized overnight at 58°C in HB containing 0.5-1 μg/ml of the corresponding riboprobe. The next day, the tissue was washed with HB for 30 minutes at 58°C. After that, we continued washing with saline sodium-citrate buffer (SSC; pH 7.5, 0.02 M), doing three washes for 15 min each at 58°C, followed by another wash in the same buffer for 15 min but at room temperature, and then one wash with Tris buffer (TB, 0.05 M, pH 7.6) for 10 min at room temperature. After, we did the inhibition of the endogenous peroxidase by incubating in 1% H_2_O_2_ and 10% methanol in TB for 30 min. After, the sections were washed with TB 0,1M three times for 10 min each. Following this, the tissue was incubated in a blocking solution (TNB) with 20% blocking reagent (BBR) and 20% of sheep serum in Tris-NaCl-Tween buffer (TNT; 10% TB, pH 8, 0.1 M; 0.9% NaCl; 0.05% Tween-20) for 2-4 h. After blocking, the tissue was incubated with a sheep anti-digoxigenin antibody diluted 1:200 (Roche Diagnostics, Basel, Switzerland) conjugated to the peroxidase enzyme, in TNB overnight at 4°C and with gentle stirring. The next day we washed with TNT three times 10 min each, and after the slices were incubated with Cy3-tyramide (AAT Bioquest) 1mM, diluted 1:200 in Tris 0,1M with H_2_O_2_ for 10 min. Finally, the sections were washed and processed for immunofluorescence against GFP.

For the immunofluorescence, the tissue was first permeabilized using PBS containing Triton100 0.3% (PBS-Tx), washed three times for 10 min each, stirring gently. After, the sections were incubated in a blocking solution with 10% normal goat serum (NGS) and 2% bovine serum albumin (BSA) in PBS-Tx 0.3% for 1-2 h at room temperature and with gentle stirring. After blocking, the tissue was incubated over weekend (from Friday afternoon until Monday morning, about 64-70 hours) with an anti-GFP primary antibody (Table 2), at 4°C while stirring gently. After the incubation, the sections were washed with PBS-Tx 0.3% three times 10 min each, and then incubated with a fluorescent secondary antibody (Table 3) in PBS-Tx 0.3% for 90-120 min at room temperature and in the dark, while stirring gently. After this, the sections were washed with PBS 0.1M for 10 min followed by Tris buffer 0.01M, rinsing two times, 10 minutes each. Finally, the sections were rinsed, mounted, and coverslipped using Vectashield Hardset Antifade mounting medium (Vector Laboratories Ltd)

**Table 2.**
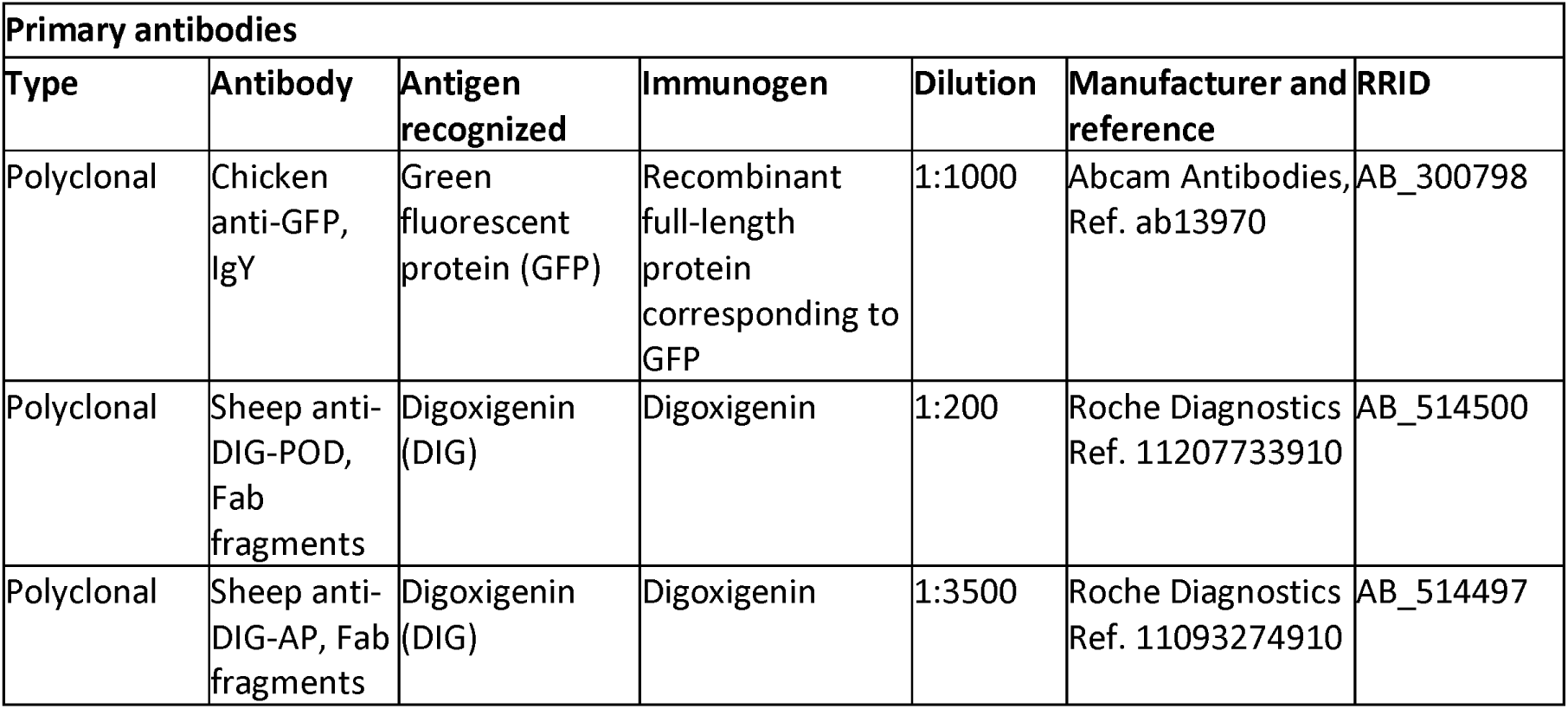
Primary antibodies.

| Primary antibodies |  |  |  |  |  |  |
| --- | --- | --- | --- | --- | --- | --- |
| Type | Antibody | Antigen recognized | Immunogen | Dilution | Manufacturer and reference | RRID |
| Polyclonal | Chicken anti-GFP, IgY | Green fluorescent protein (GFP) | Recombinant full-length protein corresponding to GFP | 1:1000 | Abcam Antibodies, Ref. ab13970 | AB_300798 |
| Polyclonal | Sheep anti-DIG-POD, Fab fragments | Digoxigenin (DIG) | Digoxigenin | 1:200 | Roche Diagnostics Ref. 11207733910 | AB_514500 |
| Polyclonal | Sheep anti-DIG-AP, Fab fragments | Digoxigenin (DIG) | Digoxigenin | 1:3500 | Roche Diagnostics Ref. 11093274910 | AB_514497 |

**Table 3.** Secondary antibodies.

| Secondary antibodies |  |  |  |  |  |
| --- | --- | --- | --- | --- | --- |
| Type | Antibody | Antigen recognized | Dilution | Manufacturer and reference | RRID |
| Biotinylated | Goat anti-chicken IgY (H + L), biotinylated | Chicken IgY | 1:200 | Vector, Ref. BA-9010 | AB_2336114 |
| Fluorescent | Goat anti-chicken IgY (H + L), coupled to Alexa 488 | Chicken IgY | 1:500 | Invitrogen, Ref. A-11039 | AB_2534096 |

### Digital photographs and figures

Digital photographs from conventional in situ hybridization sections were taken on a Leica microscope (DMR HC, Leica Microsystems GmbH) equipped with a Zeiss Axiovision Digital Camera (Carl Zeiss, Germany). Serial images from the fluorescent material were taken using a confocal microscope (Olympus FV1000; Olympus Corporation, Japan).

### Quantification

To analyze colocalization of Otp individual neurons (labeled with GFP) with *A r*, *Esr1 (ERα)*, or *Esr2 (ERβ*) mRNA, we used confocal fluorescent images of the areas of interest captured sequentially using the 40x objective. Sequential images were taken with a 2 μm-step distance between each other. Counting was done manually using the function Cell counter of Fiji ImageJ, selecting one z-level where the labelling was better for all markers and confirming the labelling with the original confocal image, using the FV10-ASW 4.2 Viewer. With this method we obtained quantitative data of the percentage of colocalization in the areas of interest.

We also used the QuPath open-source bioimage analysis software (Bankhead et al., 2017; Jerome et al., 2023) to quantify transcript signal of steroid receptors in Otp cells in confocal fluorescent images of anterior (MeA), posterodorsal (MePD) and posteroventral (MePV) subnuclei of medial amydala, posterior BSTM (BSTMp) and PVN, comparing males and females. For the QuPath analysis, we selected cases with comparable signal levels in fluorescent in situ hybridization. For *Ar*, we used images from six animals, three females and three males; for *Esr1* and *Esr2*, we used images from four animals, two females and two males. From these, we used images obtained with the 40X objectives and, for each image, we analyzed a single z level of the stack. To select the cells, we employed the cell detection tool, and then we used the subcellular detection module to analyze the number of transcripts (spots) per cell. In all cases, accurate identification was confirmed by visual inspection.

### Morphological study of the Otp neurons of the medial amygdala in both sexes

For this study, we used 6 males and 6 females of adult *Otp-eGFP* transgenic mice, aged from P271 to P337. The animals were age and litter matched.

### Sample preparation and sectioning

The fixation of the brains was done as explained previously. After this, the brains were cryoprotected with 30% sucrose in phosphate buffer saline (PBS; 0.1 M; pH 7.4) and frozen with isopentane (2-methyl butane, Sigma-Aldrich, Germany) at −60/−70°C following the protocol described by Rosene et al. (1986). The frozen brains were preserved at −80°C. The brains were sectioned in 40μm-thick frontal sections using a freezing sliding microtome (Microm HM 450; Thermo Fisher Scientific, United Kingdom). Six parallel series of free-floating sections were collected in 4°C PBS and all were processed for immunohistochemistry.

### Immunohistochemistry

The sections were permeabilized by washing with PBS containing 0.3% Triton X 100 (PBS-Tx; pH 7.4; 0.1 M), followed by an incubation with a blocking solution made with 10% normal goat serum (NGS) and 2% of bovine serum albumin (BSA) in PBS-Tx, for 1 h at room temperature. Then, the sections were incubated in the same primary antibody used for the immunofluorescence (Table 2) diluted in PBS-Tx, and the incubation was done over weekend at 4°C, while stirring gently.

After the incubation in the primary antibody, the tissue was rinsed in PBS-Tx, then processed to inhibit the endogenous peroxidase activity by an incubation in 1% H_2_O_2_ and 10% methanol in PBS during 30min, which was followed by abundant washes in PBS. Then, the sections were incubated in a biotinylated secondary antibody (Table 3) diluted in PBS-Tx for 120 min, at room temperature, while stirring gently. After this, the sections were washed with PBS-Tx and incubated with the avidin–biotin complex (AB Complex, Vector Laboratories Ltd.) for 1 h at room temperature and under gentle stirring. After, the sections were rinsed one time in PBS and then another time with Tris buffer (0.01 M, pH 8). Following this, the reaction was revealed by incubating the tissue with diaminobenzidine (DAB), diluted in a Tris-buffered solution containing urea and H_2_O_2_. Finally, the sections were rinsed and mounted using a solution of 0.25% gelatin in Tris buffer (TB; pH 8; 0.1 M), and then they were dehydrated and coverslipped with Permount (Thermo Fisher Scientific).

### Digital photographs and figures

Digital photographs were taken using a Leica microscope (DMR HC, Leica Microsystems GmbH) equipped with a Zeiss Axiovision Digital Camera (Carl Zeiss, Germany) for the images obtained with the 5x and 10x objectives, and a Zeiss microscope (Zeiss Axioskop 2) for the images obtained with the 63x objective.

### Stereological Estimations

We used digital photographs of the regions of interest captured using the 10x and 63x objectives to study if the Otp neurons of the medial extended amygdala show sexual differences regarding the volume they occupied, their cell number and their soma size using Fiji ImageJ.

For obtaining data of volume occupied by the Otp neurons inside the different nuclei of the medial extended amygdala (anteroventral, posterodorsal and posteroventral nuclei of the medial amygdala), we delimitated in all section levels of one series the regions of interest (ROIs) as the subdomains of each nucleus with a high density of Otp neurons and fibers. Using the ROI Manager function of Fiji ImageJ, we measured the ROIs areas. Volumes were estimated based on Cavalieri’s principle (Rosen & Harry 1990). The obtained volumes were corrected for brain size differences.

To analyze differences in cell number, somas of the Otp neurons inside the regions of interest were counted manually using the function Cell counter of Fiji ImageJ and the number was confirmed by checking the sections with the 40x objective using the Leica microscope.

To analyze the soma size of the neurons in the MePD we used digital photographs of the region of interest, taken at a similar level of the amygdala for all the animals and captured using the 63x objective. The size of the somas was analyzed using Fiji ImageJ to define the somas as regions of interest (ROIs) and measuring its areas with the ROI Manager function.

### Statistical analysis

The statistical analysis was done with GraphPad Prism 7, using the two-tailed Mann Whitney test. The statistical significance was stablished with p values of less than 0.05.

## Results

### Expression of sex steroid receptors in Otp neurons and circuits of the social behavior network

To understand the regulation of sexually dimorphic social behaviors, it is critical to dissect the neurons and circuits involved, and here we focused on the Otp cells of EAme and related targets in the telencephalon, hypothalamus and brainstem. We started by analyzing expression of sex steroid receptors in centers of the social behavior network containing cell bodies and/or terminals of Otp cells (Figures 1 and 2). To that aim, we processed brain sections from *Otp-eGFP* mice of both sexes for immunohistochemistry to detect GFP signal (to visualize Otp cells) and compared them with sections processed for conventional in situ hybridization to detect the mRNA of *Ar*, *Esr1 (ERα*), and *Esr2 (ERβ*) (Figs. 1, 2). In both sexes, there was expression of the three receptors genes in the different subnuclei of the EAme that also contain abundant Otp cells (mostly derived from TOH), including anterior, posterodorsal and posteroventral subdivisions of the medial amygdala, as well as ventral and posterior subdivisions of the BSTM, with differences in intensity depending on the receptor subtype and subdivision (Figure 1). Expression of *Ar* mRNA was strong in MePD (Fig. 1d), part of MePV (Fig. 1d), and posteromedial or principal subdivision of BSTM (Fig. 1h), it was moderate in the anteroventral part of MeA (Fig. 1b) and in the ventral (Fig. 1f), posterointermediate and posterolateral subdivisions of BSTM (Fig. 1h), and it was weak in the remaining parts of EAme (Fig. 1b,d,f,h). Expression of *Esr1* mRNA was moderate to strong throughout MeA, MePD, MePV and posteromedial BSTM (Fig. 1b’,d’,h’), but it was weak in other parts of EAme (Fig. 1f’). Expression of *Esr2* mRNA was moderate in the medial parts of MePD and MePV (Fig. 1d’’) and in the posteromedial BSTM (Fig. 1h’’), but weak in the rest of EAme (Fig. 1b’’,f’’,h’’).

**Figure 1:**
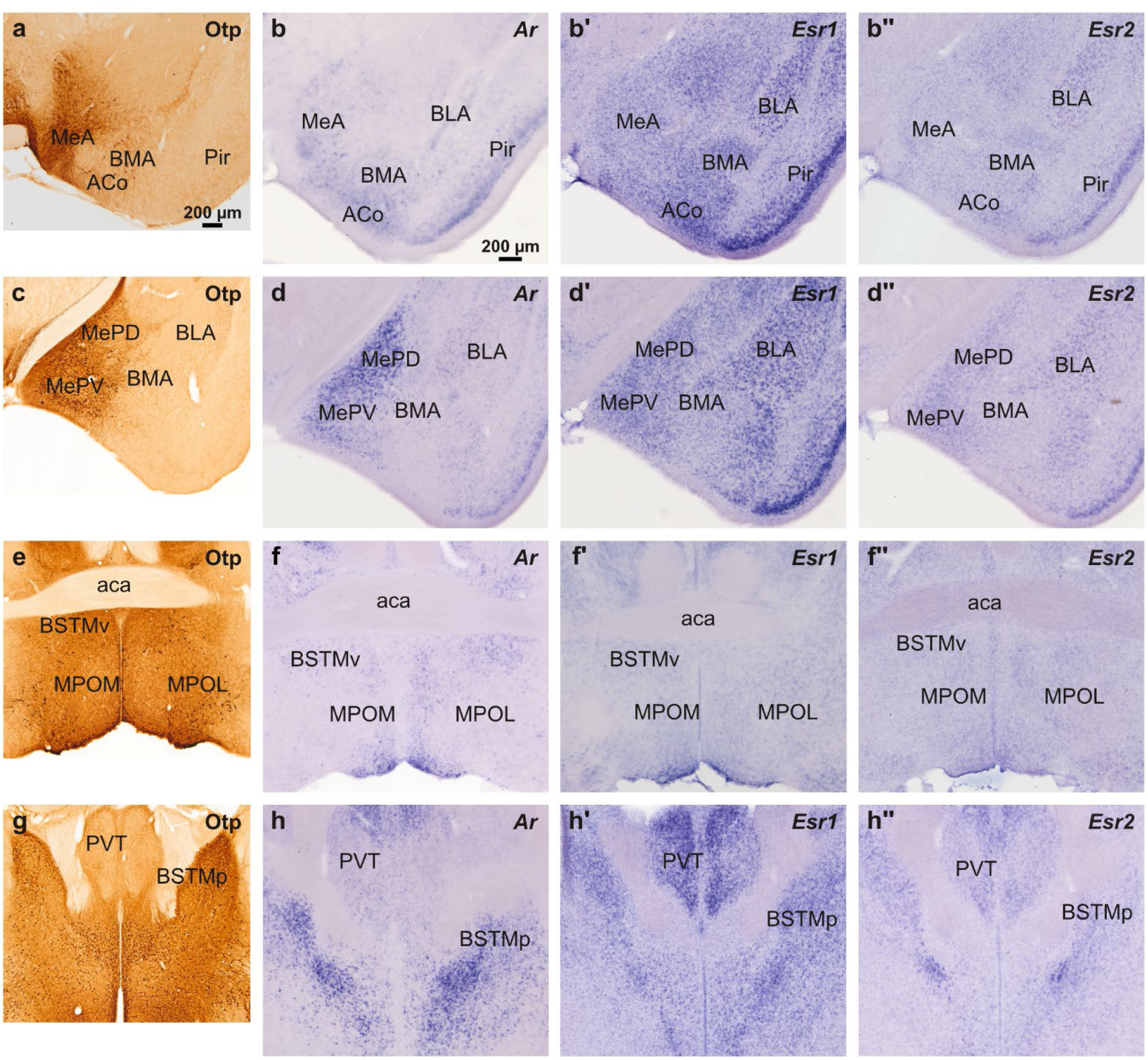
(a-h”) Images of coronal brain sections of adult Otp-eGFP mice processed for immunohistochemistry with antibodies against GFP (a, c, e, g) or for in situ hybridization to detect the mRNA of Ar (b, d, f, h), Esr1 (b’, d’, f’, h’), and Esr2 (b’’, d’’, f’’, h’’) There is expression of sex steroid receptors mRNA in areas where there are somas and neuropil of Otp (GFP immunoreactive) neurons (MeA, MePD, MePV, BSTMv, BSTMp and MPO).

**Figure 2:**
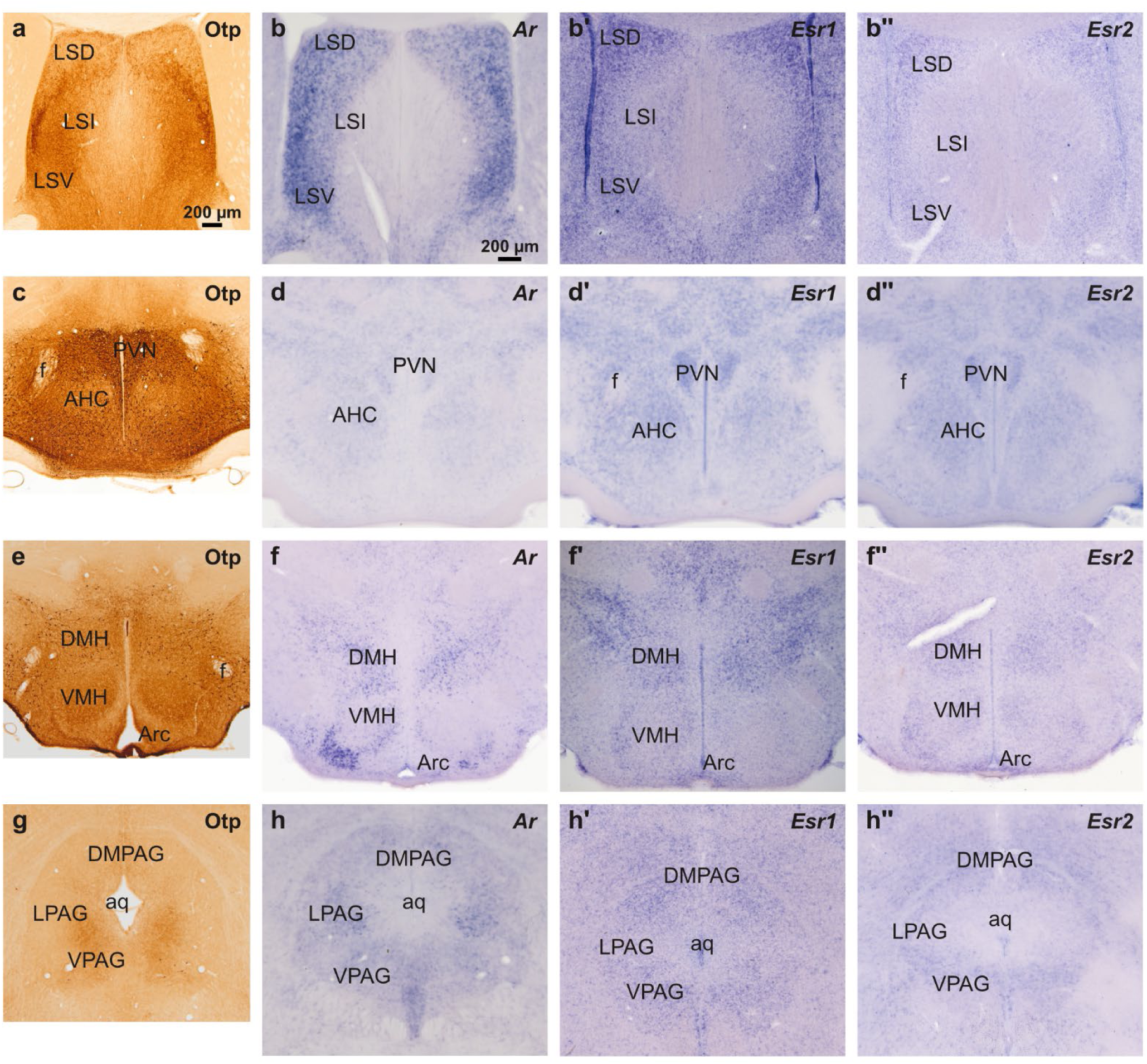
(a-h’’) Images of coronal brain sections of adult Otp-eGFP mice processed for immunohistochemistry with antibodies against GFP (a, c, e, g) or for in situ hybridization to detect the mRNA of Ar (b, d, f, h), Esr1 (b’, d’, f’, h’), and Esr2 (b’’, d’’, f’’, h’’) There is expression of sex steroid receptors mRNA in areas where there are somas and neuropil of Otp neurons (PVN, DMH and PAG) or terminals of Otp neurons (LS and VMH).

Sex steroid receptors genes were also expressed in other centers of the social behavior network, including the lateral septum (Fig. 2b-b’’), the anteroventral periventricular and medial preoptic nuclei (Fig. 1f-f’’), the PVN in the alar hypothalamus (Fig. 2d-d’’), the VMH and DMH in the basal hypothalamus (Fig. 2f-f’’), and the PAG in the midbrain (Fig. 2h-h’’). In these centers, receptor gene expression overlapped with Otp cells distribution (preoptic region, PVN, DMH, PAG) and/or axon fibers and terminals related to Otp projections (previous centers plus lateral septum and VMH). In general, expression of one or more receptors genes was more intense in the EAme nuclei regulating sexual behavior and showing sexual differences, such as MePD and BSTMpm, but also in centers known to receive heavy projections from EAme subdivisions including the lateral septum, the medial preoptic region and the ventrolateral part of VMH. Expression was particularly strong for *Ar* and *Esr1* genes in the dorsal, intermediate and ventral subdivisions of the lateral septum (Fig. 2b,b’). *Ar* and *Esr1* also showed strong expression in the anteroventral periventricular preoptic nucleus, while moderate (*Ar* and *Esr1*) or weak (*Esr2*) expression was observed in the medial part of the medial preoptic nucleus (Fig. 1f-f’’). *Ar* was also strongly expressed in the ventrolateral subdivision of VMH, while expression of estrogen receptors was weak in this nucleus (Fig. 2f-f’’). In contrast, *Ar* expression was weak or very weak in PVN, while *Esr1* and *Esr2* showed weak to moderate expression in this nucleus (Fig. 2d-d”). In addition, the three receptor genes displayed weak to moderate expression in DMH (Fig. 2f-f’’), and in PAG (with differences between its ventral, lateral and dorsal subdivisions; Fig. 2h-h’’). The lateral subdivision of PAG (receiving input from MePD and BSTMp; Canteras et al., 1995; Dong and Swanson, 2004) showed moderate expression of *Ar,* but it displayed weak expression of *Esr1* and *Esr2*.

To better understand the location of the sex steroid receptors and study if the Otp-lineage neurons of the social behavior network express these receptors, we processed brain sections of adult *Otp-eGFP* mice of both sexes for immunofluorescence against GFP combined with fluorescent in situ hybridization for each receptor. We focused on the Otp neurons of the EAme subdivisions, mostly derived from the TOH embryonic domain, and compared them with those in PVN primarily derived from the hypothalamic SPV core domain (Figs. 3-8). The results showed that in both sexes the majority of Otp neurons in all the subdivisions of EAme expressed mRNA for *Ar*, *Esr1*, and *Esr2* (between 93.5 and 100% of the Otp neurons expressed the studied receptors in all the areas of interest, Table 4). Thus, the majority of these Otp neurons are probably coexpressing those three receptors. The mRNA of the receptors was located mainly in the soma, near the nucleus or close to the cytoplasmic membrane, but there was also signal in some cell processes. In our fluorescent material, we observed variability in the expression intensity of each receptor between neurons. We observed a trend for higher percentage of *Ar* coexpression in Otp cells of MeAV, MePD and BSTMp than in those of MePV and PVN (Table 4). Moreover, in Otp cells of PVN, there was a higher percentage of coexpression with *Esr2* in females than in males (Table 4). We also observed expression of the three receptors in non-Otp cells inside the studied areas, both in the EAme subdivisions and the PVN.

**Figure 3:**
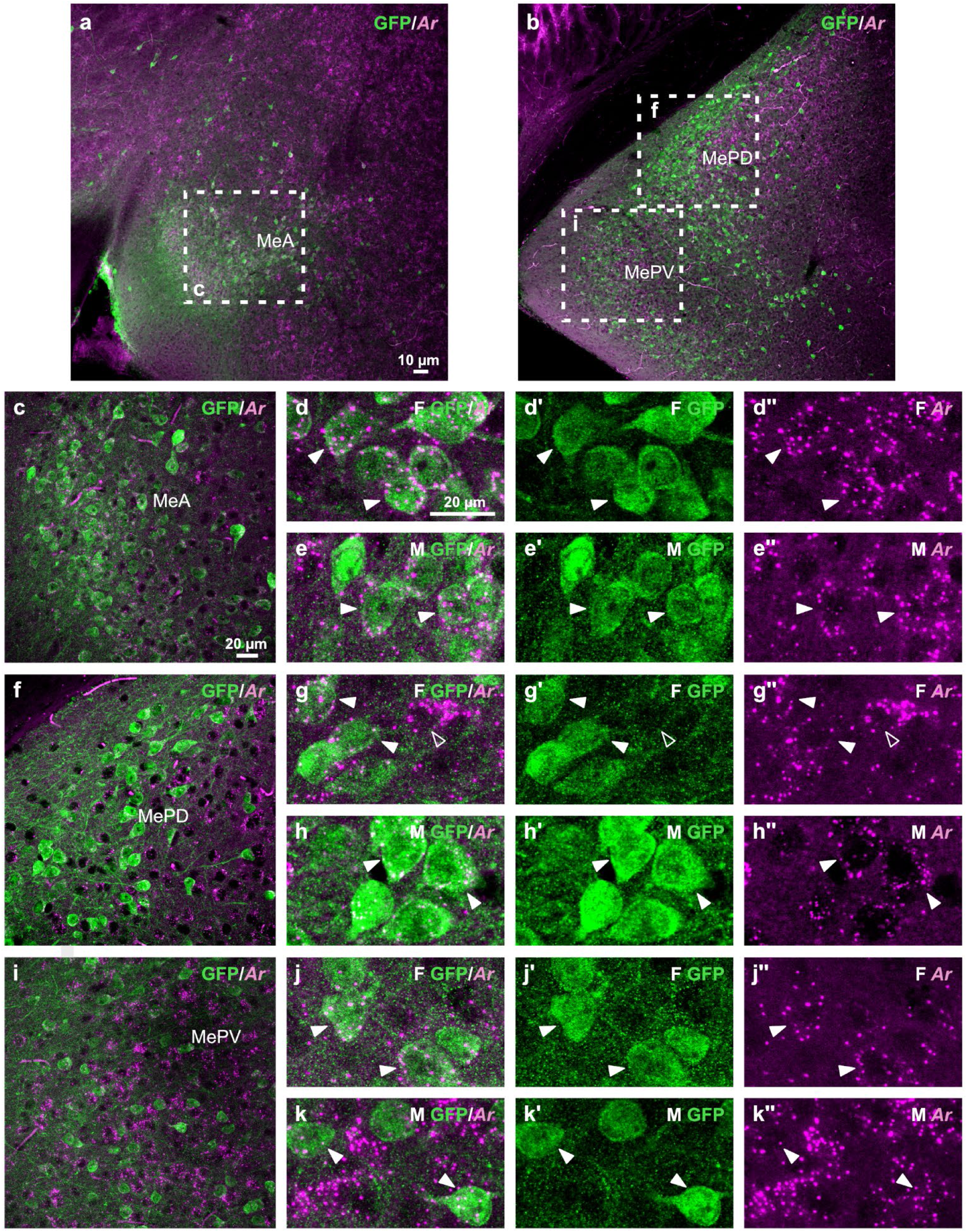
(a-k’’) Confocal images of coronal brain sections of adult Otp-eGFP mice double labeled by immunofluorescence for GFP (green) and FISH to detect Ar mRNA (magenta). The images are taken at different anteroposterior and dorsoventral levels of the medial amygdala and show that there is expression of Ar mRNA in Otp-linage neurons (filled arrowheads) and non-Otp neurons (empty arrowheads) of the MeA (c-e”), MePD (f-h”) and MePV (i-k”). The details show the expression in females (F) and males (M) (d-e’’, g-h’’, j-k’’).

**Figure 4:**
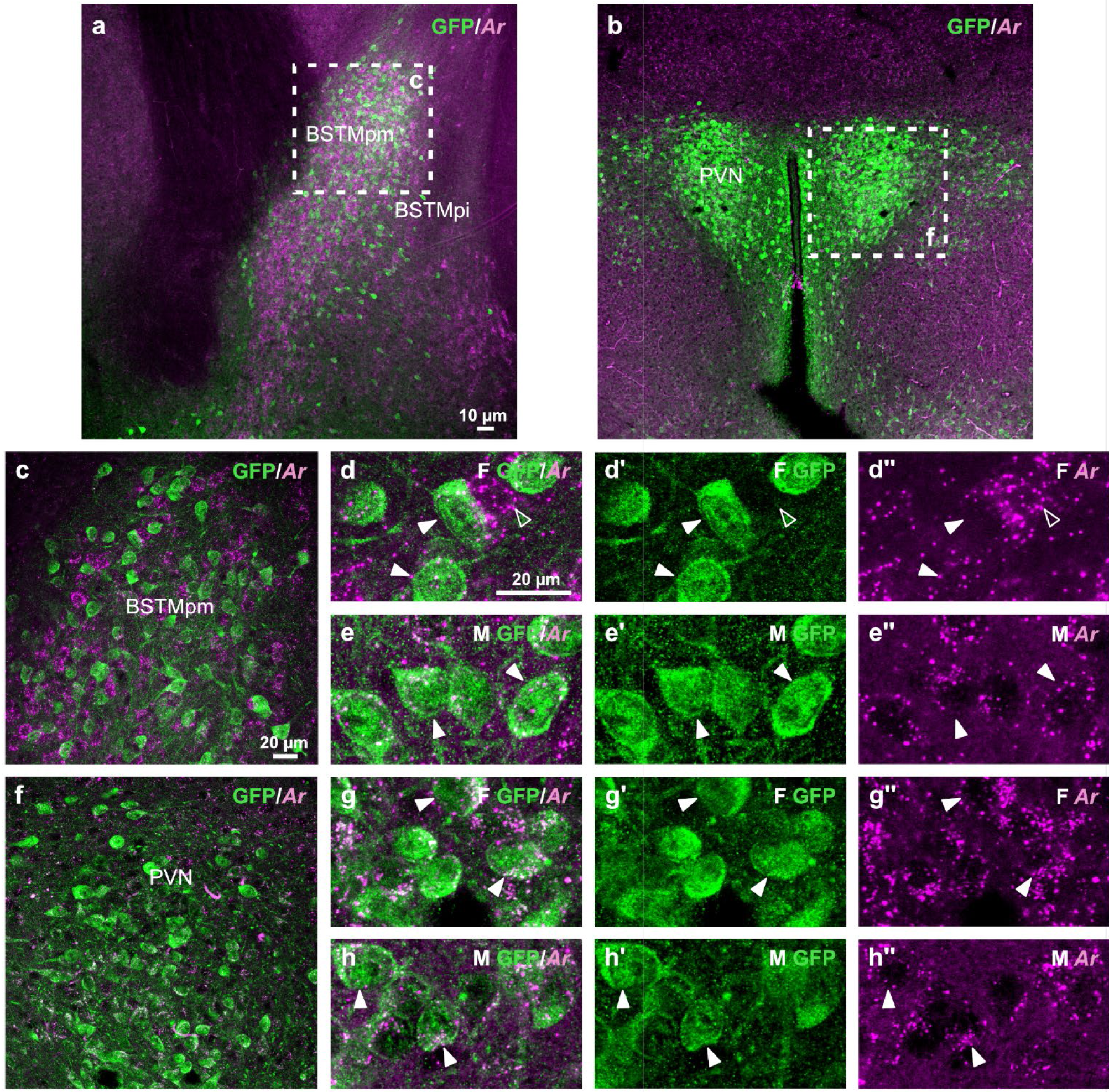
(a-h’’) Confocal images of coronal brain sections of adult Otp-eGFP mice double labelled by immunofluorescence for GFP (green) and FISH to detect Ar mRNA (magenta). The images are taken at the levels of the posteromedial BSTM (a,c, details in d-e”) and the hypothalamic PVN (b,f, details in g-h”) and show that there is expression of Ar mRNA in Otp-linage neurons (filled arrowheads) and non-Otp neurons (empty arrowheads) of the BSTM, specially BSTMpm (a, c), and the PVN (b, f). The details show the expression in females (F) and males (M) (d-e’’, g-h’’).

**Figure 5:**
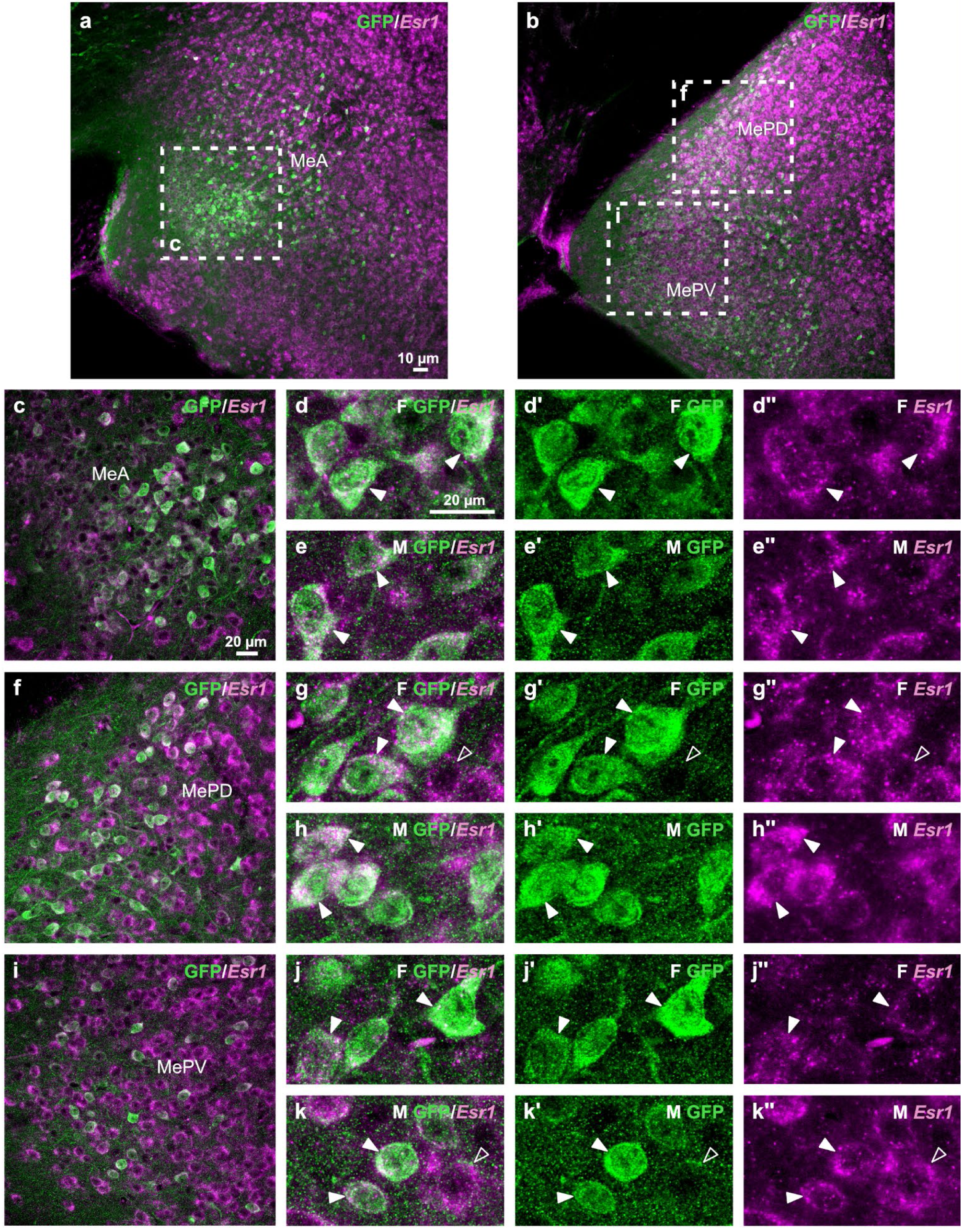
(a-k’’) Confocal images of coronal brain sections of adult Otp-eGFP mice double labeled by immunofluorescence for GFP (green) and FISH to detect Esr1 mRNA (magenta). The images are taken at different anteroposterior and dorsoventral levels of the medial amygdala and show that there is expression of Esr1 mRNA in the Otp neurons of the MeA (a,c-e”), MePD (b,f-h”) and MePV (i-k”). The details show the expression in females (F) and males (M) (d-e’’, g-h’’, j-k’’). The filled arrowheads show some examples of coexpression in Otp cells.

**Figure 6:**
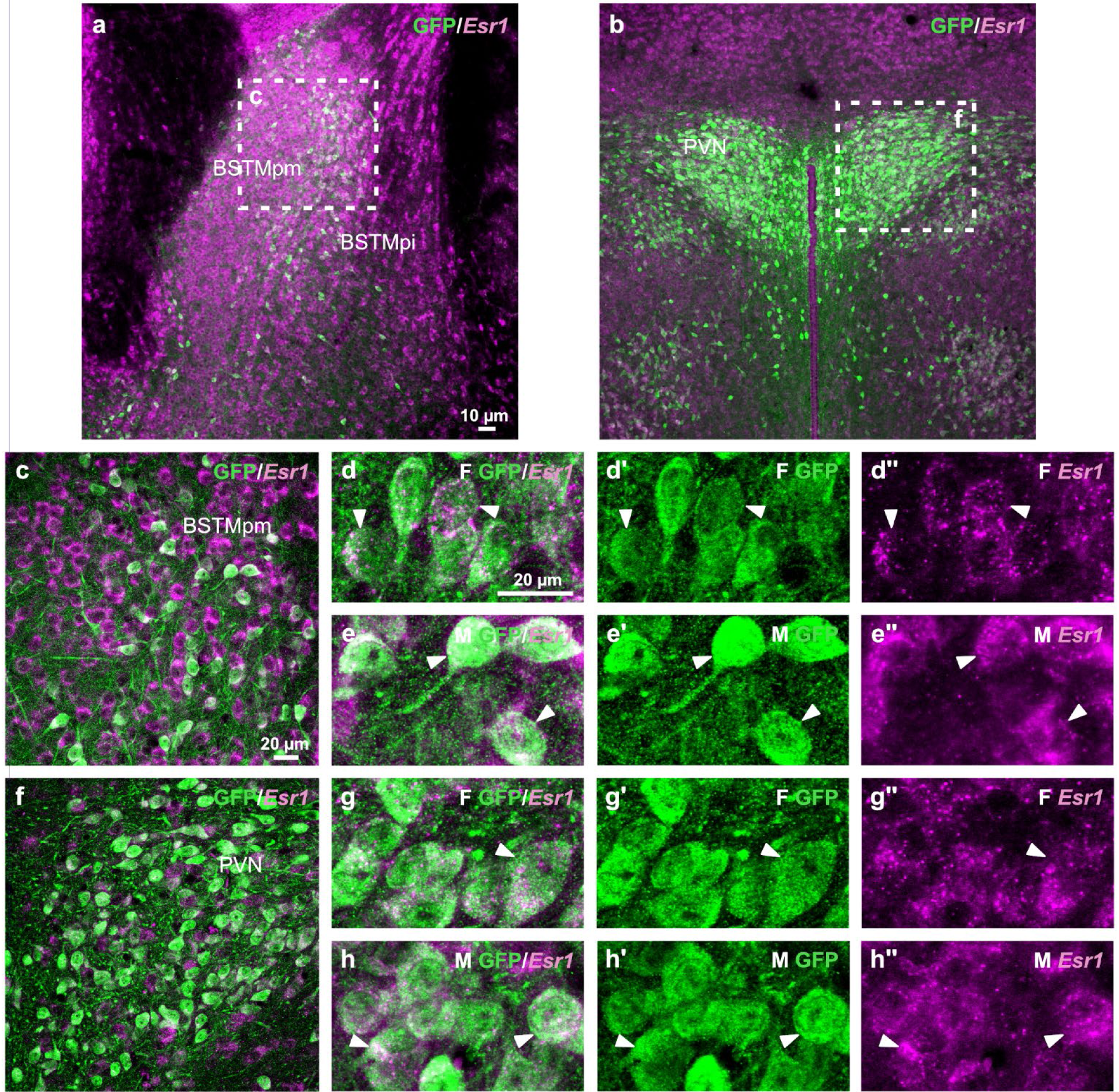
(a-h’’) Confocal images of coronal brain sections of adult Otp-eGFP mice double labeled by immunofluorescence for GFP (green) and FISH to detect Esr1 mRNA (magenta). The images are taken at the posterior BSTM (c-e”) and the hypothalamic PVN (f-h”) and show that there is expression of Esr1 mRNA in the Otp neurons of the BSTM, specially BSTMpm (a, c), and PVN. The details show the expression in the BSTMp and PVN of females (F) and males (M) (d-e’’, g-h’’). The filled arrowheads show some examples of coexpression in Otp cells.

**Figure 7:**
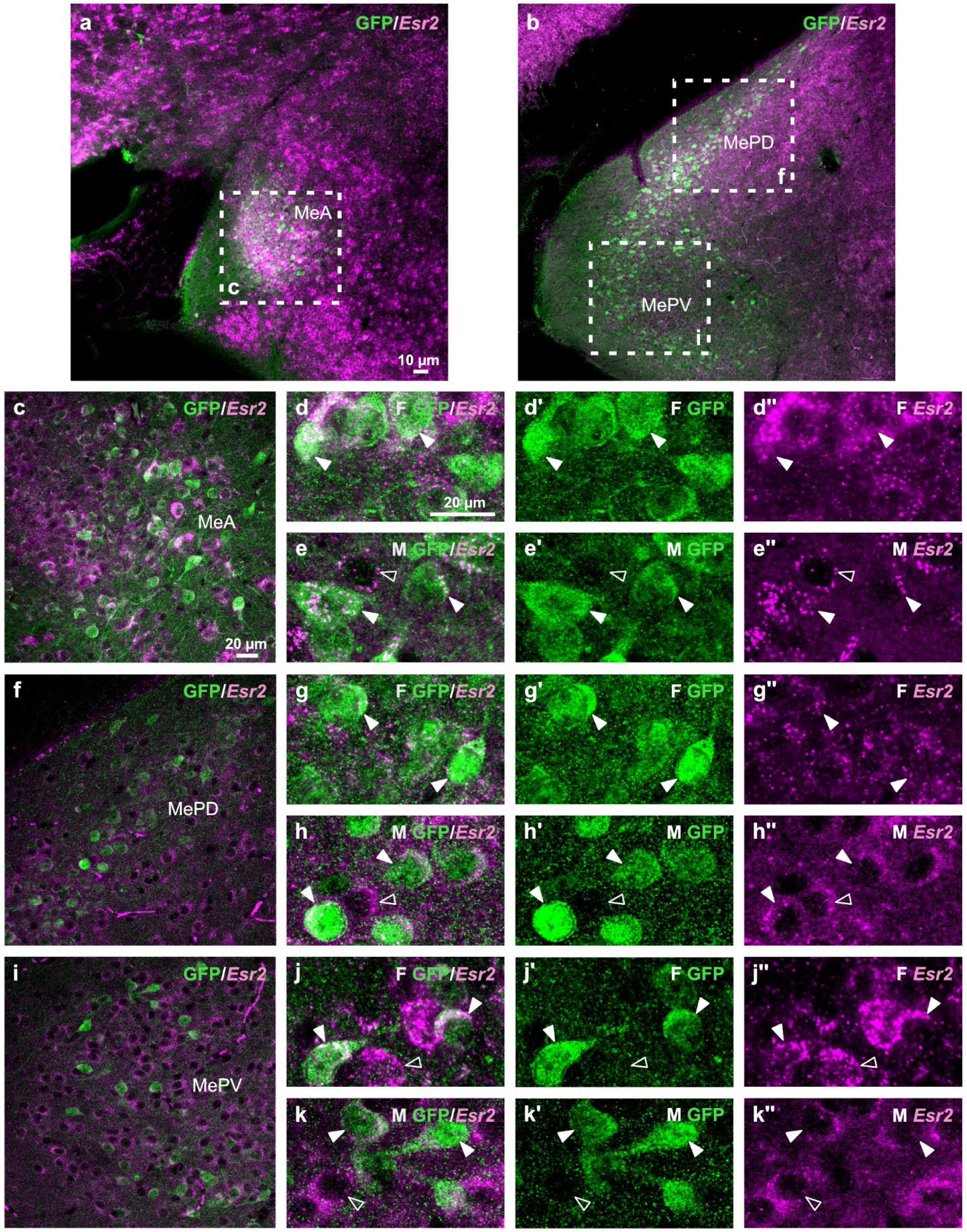
(a-k’’) Confocal images of coronal brain sections of adult Otp-eGFP mice double labeled by immunofluorescence for GFP (green) and FISH to detect Esr2 mRNA (magenta). The images are taken at different levels of the medial amygdala and show that there is expression of Esr2 mRNA in the Otp neurons of the MeA (a,c, details in d-e”), MePD (b,f, details in g-h”) and MePV (b,i, details in j-k”). The details show the expression in females (F) and males (M) (d-e’’, g-h’’, j-k’’). The filled arrowheads show some examples of coexpression in Otp cells, while the empty arrowheads show examples of expression in non-Otp cells.

**Figure 8:**
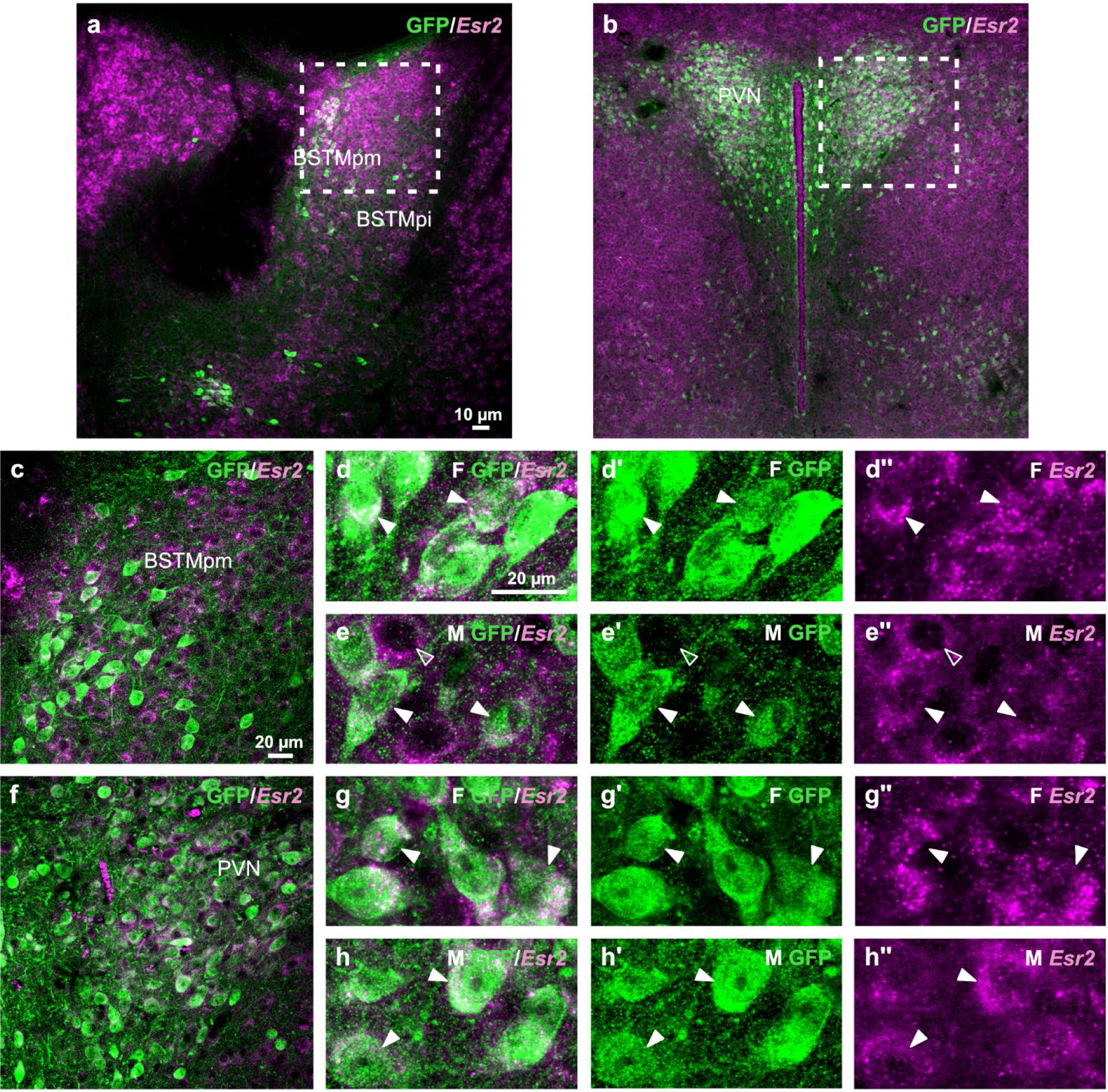
(a-h’’) Confocal images of coronal brain sections of adult Otp-eGFP mice double labeled by immunofluorescence for GFP (green) and FISH to detect Esr2 mRNA (magenta). The images are taken at the posterior BSTM (a,c-e”), and the hypothalamic PVN (b,f-h”) and show that there is expression of Esr2 mRNA in the Otp neurons of the BSTM and PVN. The details show the expression in females (F) and males (M) (d-e’’, g-h’’). The filled arrowheads show some examples of coexpression in Otp cells, while the empty arrowheads show examples of expression in non-Otp cells.

|  | Sex | Otp cells | Ar | n | Esr1 (alpha) |  |  | Esr2 (beta) |  |  |
| --- | --- | --- | --- | --- | --- | --- | --- | --- | --- | --- |
|  |  |  | coexpression |  | Otp cells | coexpression | n | Otp cells | coexpression | n |
| **MeA** | **F** | 148 | 97,3% | 3 | 93 | 98,9% | 2 | 175 | 100,0% | 2 |
|  | **M** | 138 | 100,0% | 3 | 83 | 100,0% | 2 | 239 | 99,6% | 2 |
| **MePD** | **F** | 78 | 98,7% | 3 | 74 | 98,6% | 2 | 75 | 97,3% | 2 |
|  | **M** | 103 | 99,0% | 2 | 80 | 100,0% | 2 | 109 | 100,0% | 2 |
| **MePV** | **F** | 77 | 93,5% | 2 | 118 | 100,0% | 2 | 87 | 98,9% | 2 |
|  | **M** | 74 | 94,6% | 2 | 73 | 100,0% | 2 | 109 | 100,0% | 2 |
| **BSTMpd** | **F** | 79 | 96,2% | 2 | 74 | 100,0% | 2 | 131 | 96,9% | 2 |
|  | **M** | 69 | 100,0% | 2 | 97 | 99,0% | 2 | 81 | 100,0% | 2 |
| **BSTMpv** | **F** | 58 | 98,3% | 2 | 69 | 98,6% | 2 | 80 | 96,3% | 2 |
|  | **M** | 65 | 96,9% | 3 | 74 | 97,3% | 2 | 62 | 98,4% | 2 |
| **PVN** | **F** | 248 | 96,0% | 2 | 205 | 97,1% | 2 | 208 | 99,5% | 2 |
|  | **M** | 213 | 96,2% | 2 | 163 | 98,2% | 2 | 167 | 94,6% | 2 |

To further study possible differences in expression, we analyzed the double-labeled fluorescent confocal images using QuPath. Our analysis was focused on the Otp cells of MeAV, MePD, MePV, BSTMp, and PVN. First, we selected the Otp cells by using the cell detection feature of QuPath, and the selection was confirmed by visual inspection. Then, for each nucleus and sex, we employed the subcellular detection module to obtain the number of transcripts per cell (number of spots, confirmed visually for accurate identification). For each area, we first obtained the data for each animal, and then calculated the mean per sex, plus the standard deviation (data represented in Table 5). The results showed individual differences in expression of *Ar* in Otp cells in both males and, specially, females (Table 5). In both sexes, *Ar* expression appeared to be lower in Otp cells of PVN than in those of MePD and BSTMp. With respect to estrogen receptors, *Esr1* expression in Otp cells of MePD and BSTMp subdivisions was highly variable in males (Table 5). Regarding *Esr2*, expression showed an opposite trend, but there was a high variability between individuals, specially in females. In PVN of both sexes, *Esr1* and *Esr 2*expression appeared to be higher (average number of spots per cell above 11) than *Ar* expression (average number of spots per cell below 7).

**Table 5.** QuPath analysis of *Ar, Esr1* and *Esr2* mRNA expression in Otp cells from confocal images of EAme and PVN. For *Ar*, images from six animals were analyzed, 3 females (F) and 3 males (M), while for *Esr1* and *Esr2* images from four animals were studied, 2 females and 2 males. The data (spots per cell) were obtained separately for each animal, then the mean values of all animals were calculated per each area and sex (provided in this Table). The standard deviations (SD) are included as separate columns.

|  | *Ar* |  |  |  |  |
| --- | --- | --- | --- | --- | --- |
|  | **Sex** | **Animals (n)** | **Cells (n)** | **Mean of Spots** | **SD** |
| MeA | F | 3 | 57 | 12.88 | 6.47 |
|  | M | 3 | 79 | 11.34 | 4.32 |
| MePD | F | 3 | 51 | 9.59 | 6.38 |
|  | M | 3 | 60 | 10.39 | 2.07 |
| MePV | F | 3 | 59 | 6.86 | 2.96 |
|  | M | 3 | 56 | 8.31 | 2.34 |
| BSTMp | F | 3 | 50 | 9.13 | 3.33 |
|  | M | 3 | 58 | 15.00 | 5.17 |
| PVN | F | 3 | 161 | 6.47 | 5.49 |
|  | M | 3 | 162 | 6.71 | 2.37 |

|  | ***Esr1*** |  |  |  |  |
| --- | --- | --- | --- | --- | --- |
|  | **Sex** | **Animals (n)** | **Cells (n)** | **Mean of Spots** | **SD** |
| MeA | F | 2 | 57 | 19.91 | 2.78 |
|  | M | 2 | 49 | 23.10 | 7.27 |
| MePD | F | 2 | 37 | 17.93 | 2.20 |
|  | M | 2 | 50 | 22.52 | 12.13 |
| MePV | F | 2 | 49 | 14.92 | 5.29 |
|  | M | 2 | 49 | 14.44 | 7.76 |
| BSTMp | F | 2 | 40 | 17.66 | 2.93 |
|  | M | 2 | 72 | 25.17 | 11.62 |
| PVN | F | 2 | 180 | 16.87 | 0.79 |
|  | M | 2 | 116 | 12.30 | 1.65 |
|  | ***Esr2*** |  |  |  |  |
|  | **Sex** | **Animals (n)** | **Cells (n)** | **Mean of Spots** | **SD** |
| MeA | F | 2 | 65 | 13.12 | 11.48 |
|  | M | 2 | 75 | 8.85 | 3.70 |
| MePD | F | 2 | 31 | 17.57 | 12.53 |
|  | M | 2 | 61 | 12.52 | 0.05 |
| MePV | F | 2 | 46 | 15.90 | 14.96 |
|  | M | 2 | 54 | 7.09 | 2.65 |
| BSTMp | F | 2 | 70 | 17.72 | 16.45 |
|  | M | 2 | 53 | 7.52 | 2.23 |
| PVN | F | 2 | 111 | 17.11 | 8.05 |
|  | M | 2 | 111 | 11.73 | 0.45 |

### Morphological analysis of the Otp neurons of the medial amygdala: comparison between sexes

The EAme is a nuclear complex with well-studied sexual differences in some of its subnuclei, specially MePD. Because of this, we wanted to study if the Otp neurons contribute to these differences between sexes. For this, we processed adult Otp-eGFP mice for immunohistochemistry against GFP and, by way of different ImageJ tools, analyzed different parameters of the subdomains rich in Otp neurons inside these subnuclei, such as volume, cell number and density, and the soma size.

#### Volume

For each major subnucleus of the medial amygdala, MePD, MePV and MeA, we selected the subdomain with higher density of Otp neurons and analyzed the volume occupied by these cells, comparing both sexes (Fig. 9). In MePD, the majority of the Otp neurons are located in its medial part and this subdomain was significantly bigger in males than in females after standardized by brain size (females mean = 0.084 mm^3^, SD = 0.011 mm^3^; males mean = 0.108 mm^3^, SD = 0.017 mm^3^, *U* = 2, p-value = 0.017) (Fig. 9). In contrast, we did not find differences in the subdomains rich in Otp cells of MeA and MePV. In MeA, the majority of the Otp neurons are found in the anteroventral subnucleus and there was no significant volume difference between sexes (female mean = 0.018 mm^3^, SD = 0.002 mm^3^; male mean = 0.020 mm^3^, SD = 0.003 mm^3^, *U* = 10, p-value = 0.43). In MePV, the majority of Otp neurons are found in the periphery of the subnucleus, forming a ring, while the central part only contained few Otp cells; our results showed that there were no significant volume differences in this subnucleus (female mean = 0.149 mm^3^, SD = 0.033 mm^3^; male mean = 0.149 mm^3^; SD = 0.037 mm^3^, *U* = 11, p-value = 0.54).

**Figure 9:**
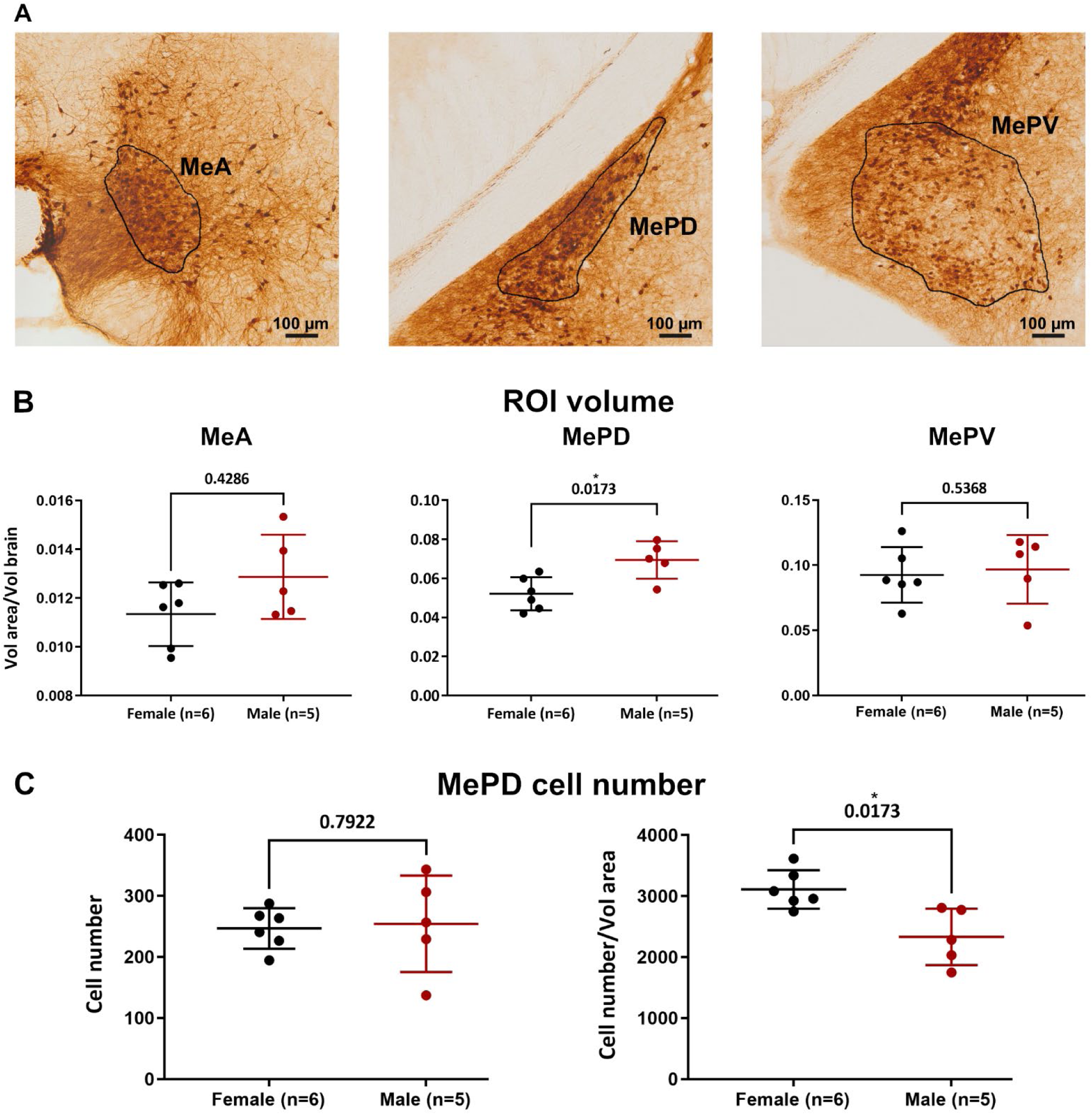
Analysis of sexual differences in the Otp neurons of the medial amygdala subnuclei. A Coronal images of sections at the level of the medial amygdala (MeA, MePD, MePV) processed for immunohistochemistry against GFP (with the 10x objective) to label Otp neurons, showing the regions of interest (ROI) delimitated for the analysis, with high density of Otp neurons. B Graphic representing the sexual differences in the volume occupied by Otp cells in subnuclei of the medial amygdala (ROI volume). Only MePD showed significant differences. C Analysis of the sexual differences in the number and density of Otp neurons in the medial subdomain of MePD. See text for more details.

#### Cell number and density

To understand better the differences observed in MePD we studied if there were differences in Otp-lineage cell number between sexes. We found that, even if the area occupied by these cells was bigger in males, the number of Otp-lineage neurons was similar between sexes (female mean = 246.7, SD= 33.3; male mean= 254.3, SD= 79.0, *U* = 13, p-value= 0.79) (Fig. 9). As a result, the density of Otp-lineage neurons, calculated dividing the number of cell bodies between the volume of the region, was significantly higher in females than in males (female mean = 3110 cells/mm^3^, SD = 315.6 cells/mm^3^; male mean = 2331 cells/mm^3^, SD = 462.7 cells/mm^3^, *U* = 2, p-value= 0.017) (Fig. 9).

#### Soma size

We also studied the soma size of the Otp-lineage neurons in MePD. We found that the soma of these neurons was significantly bigger in males than in females (female mean = 134 µm^2^, SD = 10.49 µm^2^; male mean = 149.7 µm^2^, SD = 9.6 µm^2^, *U* = 2, p-value = 0.017) (Fig. 10).

**Figure 10:**
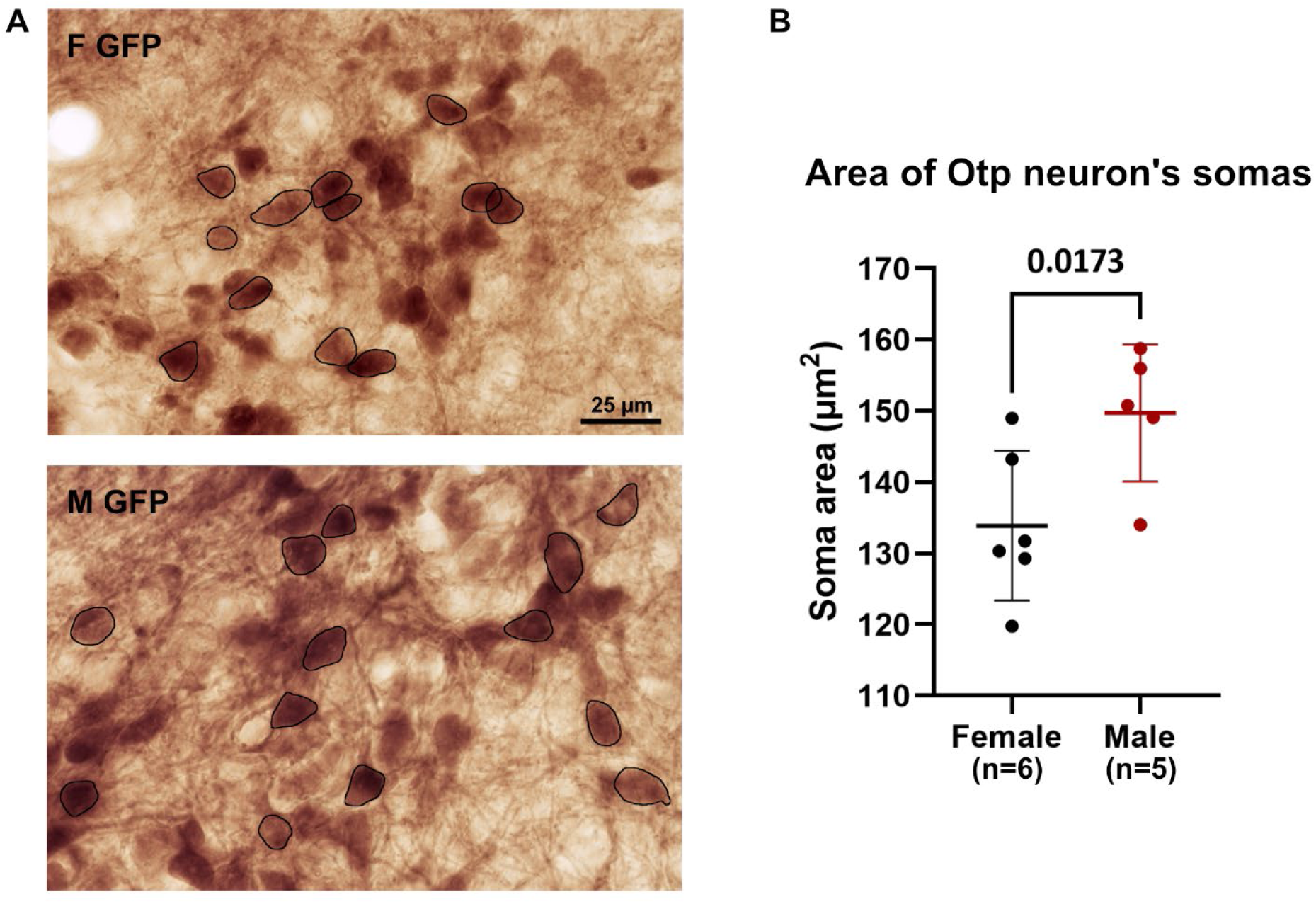
Analysis of sexual differences in the Otp neuron somas of the MePD. A Coronal images of details of Otp cells in the MePD (labeled by immunohistochemistry against GFP; 63x objective), showing a delineation of the selected somas used for analyzing the soma size in both sexes. B Graphic representing the sexual differences of the soma size of Otp neurons in the MePD.

## Discussion

The EAme is part of the social behavior network and one of the brain regions involved in sexually dimorphic behaviors in different vertebrates, including mammals and non-mammals (Newmann, 1999; De Vries and Miller, 1998; Goodson and Thomson, 2010; Xie et al., 2010; Chen and Hong, 2018; Medina et al., 2019; Han and De Vries, 2003; De Vries and Panzica, 2006). In rodents and birds, both the medial amygdala and BSTM express sex steroid receptors (Simerly et al., 1990; Shughrue et al., 1997, 1998; Lu et al., 1998; Foidart et al., 1999; Shughrue and Merchenthaler, 2001; Merchenthaler et al., 2004; Cara et al., 2021) and part of these subdivisions also show anatomical and molecular differences between sexes (De Vries and Miller, 1998; Lu et al., 1998; Cooke and Woolley, 2005; Rasia-Filho et al., 2012; Otero-García et al., 2014). The present study corroborates the mRNA expression of three sex steroid receptors, *Ar*, *Esr1* (*ERα*) and *Esr2* (*ERβ*), in the different subdivisions of EAme, as well as in EAme targets that are part of the social behavior network, such as the lateral septum, the medial preoptic area, the anterior hypothalamus, the ventromedial hypothalamic nucleus, as well as the periaqueductal gray, in agreement with previous studies (Newman, 1999; Lischinsky and Lin, 2020).

In EAme, expression is particularly high in subdivisions involved in sexual behavior, such as MePD and BSTMpm. In rats and mice of both sexes, hormone manipulations affect the size of MePD and BSTMp (Segovia and Guillamón, 1993; Morris et al., 2008). Masculinization of MePD and BSTMp is mediated by activation of both androgen and estrogen receptors, the first directly by androgens and the second by way of estradiol, after aromatization of testosterone (Cooke et al., 1999, 2003; McCarthy, 2023).

However, both the medial amygdala and BSTM are highly heterogeneous in terms of neuron subtypes, and different subpopulations are involved in regulation of different social and non-social behavioral aspects (Choi et al., 2005; Hong et al., 2014; Lischinsky et al., 2017, 2023; Johnson et al., 2021). Each subpopulation may contribute differently to the sexual differences or may not contribute at all, being involved in non-sexually dimorphic behaviors. For example, GABAergic neuron subtypes of the medial amygdala are known to promote or facilitate social interactions (Choi et al., 2005; Lischinsky et al., 2017), and at least part of them contain AR and ERα and show sexual differences (Lischinsky et al., 2017). In contrast, glutamatergic neurons appear to inhibit social interactions (Hong et al., 2014; Kwon et al., 2021) and may play a role in non-social behavior, such as defense in front of a predator (Choi et al., 2005). Although these behaviors may also be sexually dimorphic (Vinterstare et al., 2023) and the glutamatergic neurons of the medial amygdala that regulate defensive behavior also appear to modulate the activity of hypothalamic centers involved in sexual behavior (Choi et al., 2005), it was unknown whether these neurons of the medial amygdala and other parts of EAme express sex steroid receptors and are sexually dimorphic.

To further understand the role of the EAme glutamatergic neurons and their putative contribution to sexual differences, here we studied one of the major subtypes of glutamatergic neurons of EAme, those that express the OTP transcription factor during development, most of which derive from the telencephalon-opto-hypothalamic domain (TOH) and coexpress FoxG1 (Morales et al., 2021). Our results showed that the vast majority of the Otp glutamatergic neurons of the medial amygdala and BSTM in both sexes express *A r*, *Esr1* and *Esr2* mRNA. Moreover, the high percentage of receptors’ expression in the Otp neurons (between 93 and 100%) indicates that probably the majority of the Otp neurons of EAme are coexpressing the three receptors. This agrees with previous reports on coexpression of androgen and estrogen receptors in BSTM neurons of Syrian hamsters (Wood and Newman, 1995) and coexpression of both estrogen receptors in most neurons of the medial amygdala and BSTM of rats (Shughrue et al., 1998). In the Otp neurons, the mRNA of the receptors was located mainly in the neurons’ soma, near the nucleus or close to the cytoplasmic membrane, but there also was signal in the cell processes, in agreement with the cellular location reported for the receptors’ proteins (Milner et al., 2001; Mc Ewen, 2001). We also observed expression of the three receptors in non-Otp cells of EAme, which may include the GABAergic neurons of this region, such as the Foxp2+ and Dbx1-derived cells, two inhibitory neuronal lineages of the medial amygdala that express ERα and androgen receptors (Lischinsky et al., 2017). In contrast to the Otp neurons, the percentage of Dbx1-derived and Foxp2+ neurons of the medial amygdala expressing ERα or AR was lower: for ERα it ranged between 24 and 28% without differences between sexes, while for AR it ranged between 7 and 26%, with significantly higher expression in males in both neuron subpopulations (Lischinsky et al., 2017). It is likely that the receptor expression percentage is higher in the Lhx6-derived GABAergic neurons of the EAme, which are involved in sexual behavior (Choi et al., 2005), but this requires further studies.

In adult rat and mouse, previous studies showed higher expression of *Ar* mRNA and protein in BSTMpm of males (Lu et al., 1998; Shah et al., 2004), and these differences appear to develop during puberty (McAbee and DonCarlos, 1998; Cara et al., 2021). In adult mouse, the MePD also shows higher mRNA expression of *Ar* in males compared to females (Cara et al., 2021). By way of QuPath analysis of confocal images, we tried to evaluate possible differences in the expression of sex steroid receptors in the Otp cells of EAme. Our results showed a high variability between individuals in mRNA expression of *Ar, Esr1 and Esr2* in Otp glutamatergic neurons of most areas studied, including MePD and BSTMp, in both males and females.

Previous studies have shown that the level of expression depends on the species, the age, the brain area and the cell type, but it can also change between sexes and in relation to social experience and external or internal environmental conditions (for example, in the absence of gonadal hormones) (Lu et al., 1998; Równiak, 2017; Cooper et al., 2021; Cara et al., 2021). Regulation of the brain expression of sex steroid receptors is complex and dynamic, and is affected by previous experience and social status. In hamsters and Californian mice, higher AR expression in MePD has been related to social dominance in males, and higher ERα has been related to social dominance in females (Cooper et al., 2021). Moreover, disruption of *Esr1* expression in male knockout mice virtually abolished aggressive behavior (Ogawa et al., 1998). In prairie vole males, higher levels of ERα reduced prosocial behavior (Cushing et al., 2008). Its role for regulating sexual behavior appears to be predominant in females, while in males it appears to be complementary to *Ar* (Trouillet et al., 2022). Regarding *Esr2*/ERβ, previous studies in mouse and rat showed that in EAme (both MePD and BSTMp) the number of cells expressing this receptor is higher in males than in females (Morishita et al., 2023). Studies on *Esr2*-knockout male mice have shown the implication of this receptor in the establishment and maintenance of social hierarchy (Nakata et al., 2018). Further studies will be needed to investigate whether the variability between individuals in the expression of sex steroid receptors in Otp glutamatergic cells of EAme observed here is related to social status, previous experience of animals and/or other factors.

In contrast to the Otp neurons of EAme, we found that the Otp neurons of the hypothalamic paraventricular nucleus (PVN), which embryonic origin is different from most of those found in EAme (Morales et al., 2021), tend to have lower expression of *Ar* than those of EAme (mainly MeA, MePD and BSTMp). Moreover, the percentage of Otp cells of PVN coexpressing *Esr2* (ERβ) appears to be lower in males than in females (Table 4). It appears that *Esr1*/ERα and *Esr2*/ERβ play different or partially different roles during development and in the regulation of sexually dimorphic behaviors (Morishita et al., 2023). While both estrogen receptors play roles in several sexually dimorphic behaviors, studies with estrogen receptor gene knockouts and with selective agonist treatments point out that *Esr1*/ERα is essential for the development of sexually dimorphic brain features in BSTMp and other areas, while *Esr2*/ERβ is not necessary for this or play a less clear role (see discussions by McCarthy, 2023; Moroshita et al., 2023, and references within). In the PVN, which contains vasopressinergic and oxytocinergic neurons of Otp lineage (Wang and Lufkin, 2000), it appears that part of these neurons coexpress *Esr2*/ERβ (Morishita et al., 2023). Interestingly, there are species differences in the coexpression of the receptor in vasopressin or oxytocin cells, being more prevalent the first in rats, and the second in mice (Morshita et al., 2023). In addition, in the supraoptic hypothalamic nucleus (SON), which neurons originate from the same embryonic domain as those of PVN (Bardet et al., 2008; Morales et al., 2021), *Esr2*/ERβ is expressed in rats, but not mice (Morishita et al., 2023), and expression is mostly localized to the vasopressinergic neurons and less so to the oxytocinergic neurons (Kanaya et al., 2020; Morishita et al., 2023). These receptors expressed in specific Otp cells of PVN and SON may play a relevant role in sexually-dimorphic social behaviors (De Vries and Miller, 1998). As noted above, our results showed that the percentage of Otp cells of PVN coexpressing *Esr2* (ERβ) seems to be lower in males than in females, although we did not see sex differences regarding the number of transcripts (spots) per cell (variability was high). In the rat SON, the number of the *Esr2*/ERβ cells is also higher in females than in males, but this is only seen when analyzing the protein, but not the mRNA (Morishita et al., 2023). This is an important observation, as our study is based on mRNA, but not protein expression. Sex (and perhaps species) differences in *Esr2*/ERβ expression may be partly caused by post-transcriptional and/or post-translational regulation (Morishita et al., 2023). Expression levels of this receptor also fluctuate depending on the circulating estrogens (Morishita et al., 2023), which could have a potential impact on the results of expression studies and comparisons between species, ages and sexes.

Overall, our results on the expression of sex steroid receptor mRNA by Otp glutamatergic neurons of EAme and PVN, together with previous data on other neuronal subpopulations of EAme (Lischinsky et al., 2017), highlight that both GABAergic and glutamatergic neurons of EAme express sex-steroid receptors in area and cell-type-specific manners and that embryonic origin is an important determinant for the level of expression. Although additional factors are involved (as discussed by Cara et al., 2021; and Morishita et al., 2023), the reported differences between areas and sexes in the expression of some of the sex-steroid receptors could contribute to shape differences in sex-specific behaviors (for example, sex differences in stress vulnerability in relation to social status; Cooper et al., 2021), but these could also depend on the neuronal subtype involved, as different neurons regulate different aspects of behavior (Choi et al., 2005; Hong et al., 2014; Lischinsky et al., 2017, 2023). In relation to this, microinjections of either glutamate or GABA in MePD of awake rats are known to change cardiovascular baroreflexes in opposite directions, with glutamate activating the sympathetic system and associated cardiac and vascular modulations, and GABA activating the parasympathetic systems and related effect on decreasing heart rate (Neckel et al., 2012). It is unknown whether activation of either glutamatergic and GABAergic neurons of MePD and other parts of EAme, some of them with higher expression of Ar in males than in females, would have similar effects to those just mentioned (which could be mediated by local connections, as well as by projections to autonomic-related centers in the hypothalamus and brainstem), and additional studies would be needed.

As mentioned above, the MePD is a sexually-dimorphic nucleus, well known for being androgen-responsive and for regulating sexual behavior specially in males (Cooke, 2005; Morris et al., 2008; Rasia-Filho et al., 2012; Zancan et al., 2017). In rats and mice, the differences are seen in volume, soma size and density of dendritic spines, all of which are higher in males, and these differences are lost following castration (Morris et al., 2008; Rasia-Filho et al., 2012; Pfau et al., 2016; Zancan et al., 2017). These studies did not consider the different subpopulations of MePD neurons, which include several GABAergic and glutamatergic subtypes. Since the Otp glutamatergic neurons of MePD show expression of sex steroid receptors (present results), we aimed to analyze if these specific neurons also displayed differences in terms of volume occupied, density, cell number, and soma size. Because Otp glutamatergic cells are also found in other subnuclei of the medial amygdala, we carried out the study in the anteroventral (MeAV), MePD and MePV subdivisions. We observed sexual differences in the volume occupied by the Otp neurons inside the MePD, as well as in their soma size, being significantly larger in males than in females. However, we found no differences in Otp cell number, but the Otp cell density was significantly higher in females. Previous studies have related the volume differences to more complex dendritic arbors and spines, as well as denser axon terminals in MePD of males (Rasia-Filho et al., 2012). The larger Otp neurons of male MePD found in our study may also have more complex dendritic arbors and connections, although our material did not allow to analyze dendritic branching and spine density. Overall, our results are consistent with those already described in mice by analyzing the nucleus as a whole (Cooke, 2006; Morris et al., 2008; Rasia-Filho et al., 2012), and point to the Otp neurons as major contributors to the sexual differences of MePD. In contrast, we found no differences between sexes in the Otp neurons of the MeA and MePV, at least regarding the volume occupied by these neurons, which is in line with previous studies considering the whole subnuclei. Nevertheless, since volume delineation of Otp cells of MeA and MePV is tricky because they are not as compact as those of MePD, more studies are necessary to know if there are sexual differences in the Otp cells of MeAV and MePV in terms of soma size and dendritic complexity. This should also be done for the Otp neurons of the sexually-dimorphic BSTMpm subnucleus.

In summary, our results showed that the vast majority of Otp glutamatergic neurons of MePD and BSTMpm express mRNA of *Ar*, *Esr1* and *Esr2* receptors in both males and females. Moreover, Otp neurons are larger, occupy more space and possibly possess more complex dendritic arbors in males than in females. These differences likely contribute to some of the sexually-dimorphic behaviors regulated by EAme, and could also be mediating sex-specific responses to psychosocial stressors. In this context, it would be interesting to study the expression of these receptors in the Otp cells of MePD and BSTMp at critical stages of amygdala development. Moreover, considering the abundance of this subpopulation of glutamatergic neurons in EAme, it would be interesting to study their implication in different behaviors.

## Ethical Statement

We declare that we adhere to the ethical and integrity policies of the journal regarding research.

## Availability of data

The most relevant data that support the findings of this study are included in the Figures of this paper.

## Acknowledgments

We deeply thank all Agencies that funded our research. We also thank the technicians and other staff of the Department of Experimental Medicine, as well as the Rodent Animal Facility of the University of Lleida, the Service of Proteomics and Genomics of the University of Lleida, the Service of Statistics of IRBLleida, and the Service of Genomics of the University of Valencia.

## Author contributions

ED and LMed first designed the project, and AGA contributed to refine it. AGA processed most of the material as part of her Ph.D. research project, and LMor contributed in part of the processing. AGA photographed, analyzed and prepared the figures, and analyzed the material with help of LMed and ED. ED supervised and revised the analysis on cell counting. ES helped with the QuPath analysis and the interpretation of the data. AGA produced the first draft of the manuscript, LMed revised it, ED contributed to improve it, and all authors revised it and approved it.

## List of abbreviations

aca: anterior commissure
ACo: anterior cortical amygdalar area
Arc: arcuate nucleus
BLA: basolateral amygdalar nucleus
BMA: basomedial amygdalar nucleus
BSTM: medial bed nucleus of the stria terminalis
BSTMv, BSTM: ventral subucleus
BSTMp, BSTM: posterior part
BSTMpm, BSTM: posteromedial subnucleus
DMH: dorsomedial hypothalamic nucleus
f: fornix
LS: lateral septum
MeA: medial amygdala, anterior part
MeAV: medial amygdala, anteroventral subnucleus
MeP: medial amygdala, posterior part
MePD: medial amygdala, posterodorsal subnucleus
MePV: medial amygdala, posteroventral subnucleus
MPO: medial preoptic nucleus
PAG: periaqueductal gray
Pir: piriform cortex
PVN: paraventricular hypothalamic nucleus
PVT: paraventricular thalamic nucleus
VMH: ventromedial hypothalamic nucleus

